# The ancient fusogen EnvP(b)1 is expressed in human tissues and its structure informs the evolution of gammaretrovirus envelope proteins

**DOI:** 10.1101/2020.04.22.056234

**Authors:** Kevin R. McCarthy, Joseph L. Timpona, Simon Jenni, Vesna Brusic, Welkin E. Johnson, Sean P.J. Whelan, Lindsey R. Robinson-McCarthy

## Abstract

Host genomes have acquired diversity from viruses through the capture of viral elements, often from endogenous retroviruses (ERVs). These viral elements contribute new transcriptional control elements and new protein encoding genes, and their refinement through evolution can generate novel physiological functions for the host. EnvP(b)1 is an endogenous retroviral envelope gene found in human and other primate genomes. We show that EnvP(b)1 arose very early in the evolution of primates, i.e. at least 40-47 million years ago, but has nevertheless retained its ability to fuse primate cells. We have detected similar sequences in the genome of a lemur species, suggesting that a progenitor virus may have circulated 55+ million years ago. We demonstrate that EnvP(b)1 protein is expressed in multiple human tissues and is fully processed, rendering it competent to fuse cells. This activated fusogen is expressed in multiple healthy human tissues and is under purifying selection, suggesting that its expression is selectively advantageous. We determined a structure of the inferred receptor binding domain of human EnvP(b)1, revealing close structural similarities between this Env protein and those of currently circulating leukemia viruses, despite poor sequence conservation. This observation highlights a common scaffold from which novel receptor binding specificities have evolved. The evolutionary plasticity of this domain may underlie the diversity of related Envs in circulating viruses and coopted elements alike. The function of EnvP(b)1 in primates remains unknown.

**SIGNIFICANCE STATEMENT:** Organisms can access genetic and functional novelty by capturing viral elements within their genomes, where they can evolve to drive new cellular or organismal processes. We demonstrate that a retrovirus envelope gene, EnvP(b)1, has been maintained as a functional protein for 40 to ≥55 million years and is expressed as a protein in multiple healthy human tissues. We believe it has an unknown function in primates. We determined the structure of its inferred receptor binding domain and compared it with the same domain in modern viruses. We find a common conserved architecture that underlies the varied receptor binding activity of divergent Env genes. The modularity and versatility of this domain may underpin the evolutionary success of this clade of fusogens.

## INTRODUCTION

Integration into nuclear DNA is an obligate step in the retrovirus replicative cycle. Integration events in gamete cells or their progenitor cells can allow viral elements to be vertically transmitted to offspring. Approximately 8% of the human genome is derived from such endogenous retroviruses (ERVs), and similar percentages are observed across the animal kingdom (1). Most ERV loci have acquired inactivating mutations that erode the coding potential of individual genes and render these loci incapable of producing infectious viruses. However, some ERV genes can retain their function in an otherwise degraded provirus, an indication that selective pressures preserve the function of these genes. Over time, ERV genes can be coopted to perform beneficial or essential host functions. For example, the cell-cell fusion events that produce the placental syncytiotrophoblast layer in eutherian mammals are driven by multiple independently acquired and independently coopted ERV envelopes (Envs), known as syncytins (2). Primate genomes harbor a number of ERV Env open reading frames (ORFs) with unknown functions (3, 4). Many of these ORFs have been maintained for the duration of primate evolution, time periods exceeding 40 million years.

The human genome contains four copies of ERV-P(b), located on different chromosomes; one of these, located within an intron of the *RIN3* gene, retains an intact ORF termed EnvP(b)1 (5). The intact EnvP(b)1 ORF has been identified in Old World primates and must have arisen as the result of integration of a then-extant virus into the *RIN3* locus of a common primate ancestor. The primate EnvP(b)1 ORF is reportedly evolving under purifying selection, and various dating methods estimate EnvP(b)1 to be between 30-55 million years old (5, 6). EnvP(b)1 has the characteristics of a classical gammaretroviral envelope protein. All the sequence motifs necessary for processing and cell surface expression remain intact (5, 7). When ectopically expressed, human EnvP(b)1 can mediate fusion of cells in culture (7, 8). EnvP(b)1 RNA has been detected in human tissues; however, protein expression and processing have not been previously reported (5, 6). It is unclear what function(s) EnvP(b)1 may have that explain the fact that it has maintained a functional ORF over tens of millions of years.

Here, we show that EnvP(b)1 protein is expressed in multiple human tissues in a form that appears to be fusion competent. We also demonstrate that EnvP(b)1 is more ancient than previously reported by identifying orthologs in apes, Old World monkeys and New World monkeys (simian species). These findings indicate that this element was acquired no more recently than 40-47 million years ago (MYA). Identification of similar sequences in the genome of a lemur species suggests that an infectious progenitor virus may have circulated over 55 MYA. The activity of EnvP(b)1 as a membrane fusogen has also been preserved for the duration of simian evolution. The host factors required for fusion appear to be conserved and ubiquitously expressed in the cultured cells of primates. Thus, an activated, ancient fusogen of unknown function is expressed in multiple healthy human tissues. We determined the structure of the EnvP(b)1 inferred receptor binding domain and compared it to structures of the homologous domain from two extant retroviruses. The similarity of these structures, apparent even after tens of millions of years of divergence and selection for different functions, highlights the flexibility of the architecture of this domain as a versatile scaffold for receptor binding.

## RESULTS

### EnvP(b)1 protein is expressed in human tissues

EnvP(b)1 is the only remaining intact ORF in the ERV-P(b) provirus in the human *RIN3* locus, ERV-P(b)1. All other ERV genes at this locus have been disrupted. The phylogenetic tree in Fig. 1A describes the evolutionary relationships of EnvP(b)1 with other retroviral Envs. EnvP(b)1 is part of a clade of viruses that comprises endogenous mammalian and extant avian examples. Based upon its phylogenetic relationships with other Envs and characteristic motifs in its surface (SU) and transmembrane (TM) domains, EnvP(b)1 has been previously classified as a gamma-type Env (5). Notable elements include an intact furin cleavage site and a CXXC/CX_6_CC disulfide bond motif (Fig. 1B).

**Fig. 1.**
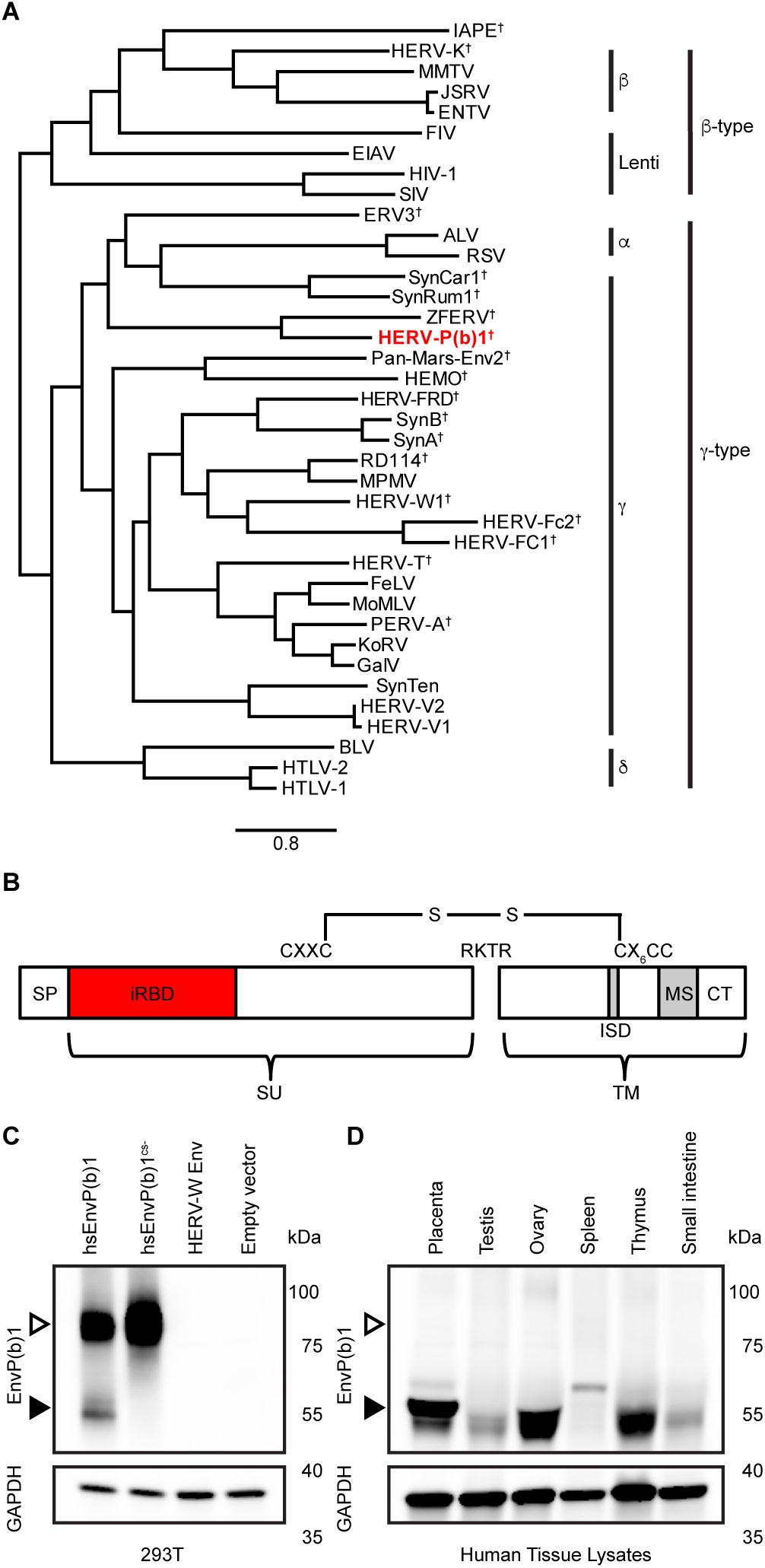
EnvP(b)1 protein is expressed and processed in human tissues (A) Maximum likelihood tree of retroviral Envs. Full-length amino acid sequences of Envs from five retroviral genera were used as inputs. The tree is rooted on IAPE Env. Genera of Env are indicated, as well as the type of Env based on phenotype and sequence (beta-type and gamma-type). † indicates sequences that have only been found as endogenous sequences. EnvP(b)1 from human ERV-P(b) (HERV-P(b)1) is shown in red. Scale bar: amino acid substitutions per site. (B) Diagram of mature EnvP(b)1 protein. SU: surface subunit. TM: transmembrane subunit. SP: signal peptide. iRBD: inferred receptor binding domain. ISD: immunosuppressive domain. MS: membrane spanning domain. CT: cytoplasmic tail. Furin cleavage site (RKTR) is noted. C. Lysates from 293T cells transfected with the indicated envelope were subjected to western blotting with either anti-EnvP(b)1 sera or anti-GAPDH antibody. Open arrows indicate the molecular weight of unprocessed SU-TM. Filled arrows indicate processed SU. D. Expression of EnvP(b)1 protein in healthy human tissues. Protein lysates from the indicated tissues from healthy donors were subjected to western blotting with either anti-EnvP(b)1 sera or anti-GAPDH antibody. A single band corresponding to EnvP(b)1 SU is present in all tissues except spleen.

EnvP(b)1 RNA has previously been detected in human tissues (5, 7). To determine whether EnvP(b)1 protein is also present, we expressed the genetically unique (no similar sequences in human or rabbit genomes) inferred receptor binding domain (iRBD) and used this iRBD protein to immunize a rabbit and produce antisera. We enriched for EnvP(b)1 iRBD-binding IgGs with a series of affinity resins and confirmed by western blot that the enriched iRBD antisera detected EnvP(b)1 in transfected 293T cells (Fig. 1C). Two major bands were observed, one corresponding to the predicted molecular weight of unprocessed EnvP(b)1 SU-TM and a second corresponding to the furin-processed SU domain. The specificity of the SU band was confirmed by expressing EnvP(b)1 in which the furin cleavage site was mutated (EnvP(b)1^CS-^). For this mutant, only the band corresponding to the unprocessed Env was observed. The reagent is specific for EnvP(b)1, as it failed to detect human Syncytin-1 (HERV-W Env), and western blots of mock transfected 293T lysate did not show bands in the appropriate locations.

To determine whether EnvP(b)1 protein is expressed in healthy human tissues, we assayed tissue lysates by western blot (Fig. 1D). We chose tissues that were similar to those in which RNA transcripts were previously detected (5, 7). Protein expression was strongest in placenta, ovary, and thymus. In these tissues we detected a doublet band for SU that was most prominent in placenta. This may represent an additional proteolytic processing event or glycosylation heterogeneity. EnvP(b)1 protein was also detected at lower levels in testis and small intestine. A band of a different molecular weight was present in spleen lysate, which we therefore cannot conclude corresponds to EnvP(b)1 protein. Tissues from both healthy males and females showed clear EnvP(b)1 expression, indicating that expression is not restricted to females (Table S1). We did not observe a higher molecular weight species corresponding to unprocessed SU-TM in any tissue, suggesting that EnvP(b)1 is efficiently processed in these tissues. In contrast, overexpression of EnvP(b)1 in 293T cells produced primarily SU-TM (Fig. 1C).

### EnvP(b)1 ORF is conserved in simian primates

The EnvP(b)1 ORF has been identified previously in select ape and Old World monkey genomes (5). We surveyed publicly available databases of mammalian, avian, reptilian, amphibian, piscine and insect genomes for EnvP(b)1 related sequences. Our survey identified the intact EnvP(b)1 ORF in 35 simian species (New and Old World monkeys and apes) We verified that the 35 simian examples were present within the *RIN3* locus by using assembled genomes and examining sequences that flanked the EnvP(b)1 ORF. The phylogenetic relationships between the simian EnvP(b)1s generally follow the same clustering as their hosts (Fig. 2A). This pattern of branching is consistent with insertion of an ancestral ERVP(b) into the *RIN3* locus of a common ancestor of all extant simian primates, followed by divergence of the Env sequences together with their respective host species. Based on the reported divergence times of New World and Old World monkey lineages (9–11), the EnvP(b)1 insertion arose at least 40-47 MYA – considerably earlier than previous synteny-based estimates (5).

**Fig. 2.**
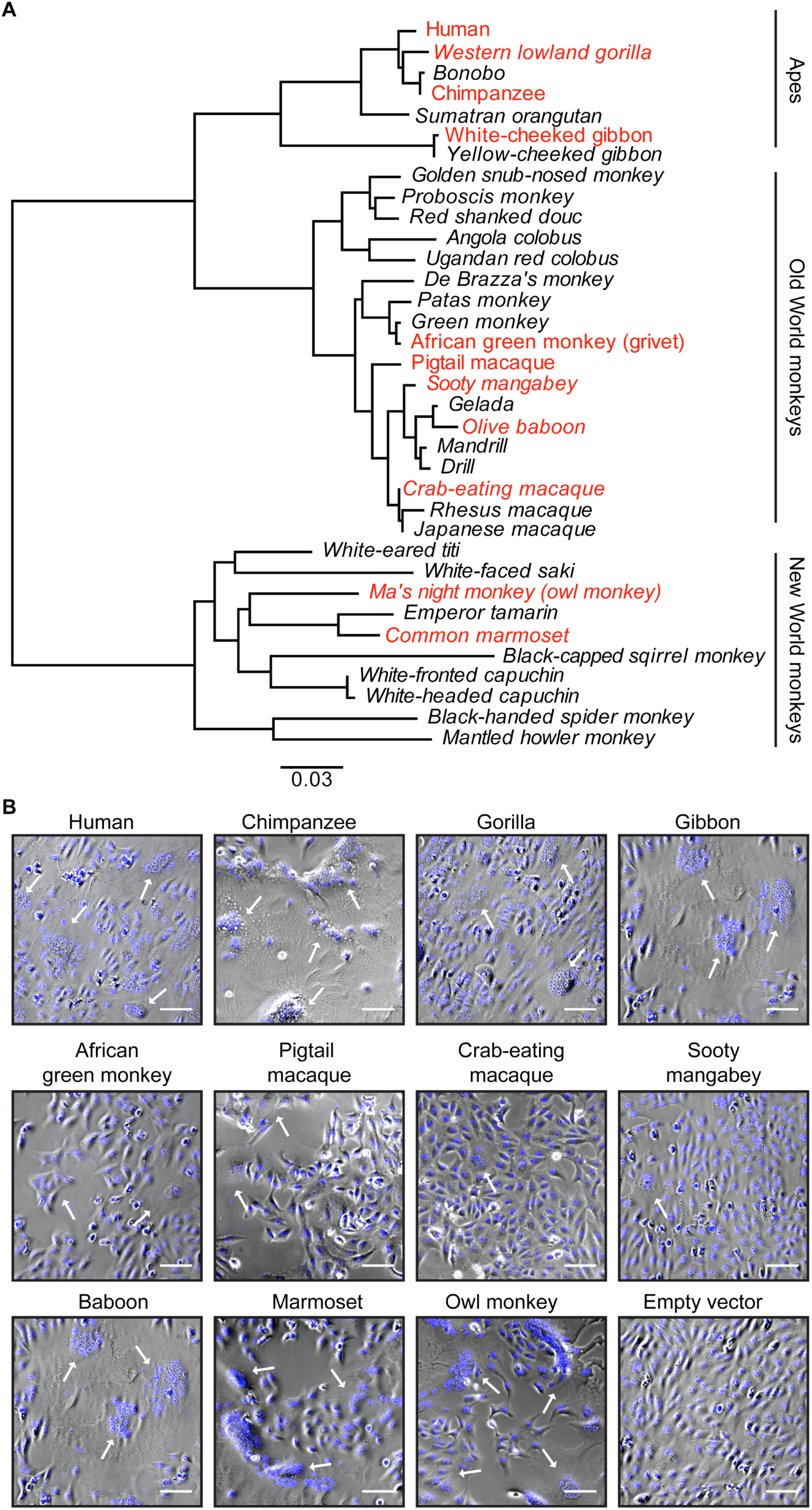
EnvP(b)1 is a conserved fusogen of simian primates. (A) Maximum likelihood tree of all full-length amino acid sequences of EnvP(b)1 ORFs identified from publically available databases. Sequences in italics are newly identified in this manuscript. Sequences highlighted in red were used for further experiments. (B) Examples of cell fusion results. Vero (green monkey) cells were transfected with EnvP(b)1 ORFs from the indicated species. Cells were stained with nuclear stain (blue) and imaged. Arrows indicate multinucleated syncytia arising from EnvP(b)1-mediated cell fusion. Scale bar: 100 µm.

Our survey of genomes identified a number of sequence reads from an incomplete aye-aye (lemur) genome. These reads are related to EnvP(b)1 and other proviral sequences including the ERV-P(b) LTRs (Fig. S1A). Overlapping, but unique, sequences suggest that at least two ERV-P(b)-related loci are present in aye-aye genomes. At least one copy has lost its Env coding potential. Two non-overlapping reads cover 69% of the iRBD. How these sequences entered the aye-aye genome is unknown. Lemurs became geographically isolated on Madagascar between 55-75 MYA (12, 13). Aye-ayes, which occupy a basal branch of lemurs, diverged from other extant lemur populations ~55 MYA (9–11, 14). Three potential scenarios for ERV-P(b) endogenization are outlined in Fig. S1C.

### EnvP(b)1 orthologs fuse cells of diverse tissue and species origin

EnvP(b)1 has evolved under at least three consecutive stages comprising distinctly different selective pressures: 1) as the Env protein of a circulating virus previous to endogenization, 40-47 MYA; 2) as a captured ERV during the process of cooption by the host; and 3) as a gene under purifying selection in all simian species. The fact that the EnvP(b)1 ORF has been retained throughout simian evolution suggests that it may perform a host function(s). Expression of human EnvP(b)1 has been shown to drive the formation of multi-nucleated syncytia in cultured cells, an indication of cell-cell fusion (7, 8). We examined the expression, proteolytic processing and ability to mediate cell-cell fusion of 11 divergent simian EnvP(b)1 orthologs that were expressed from cells of human, Old World monkey and New World monkey origin. All EnvP(b)1s were expressed in all cell lines, as shown by western blot (Fig. S2). Similar to our observations in 293T cells, and unlike human tissue lysate, the dominant form of the protein was unprocessed SU-TM. EnvP(b)1 orthologs from New World monkeys (marmoset and owl monkey) were detected at lower levels than those from apes and Old World monkeys, possibly due to sequence differences in the iRBD compared to the human ortholog (against which the sera was raised). All 11 EnvP(b)1 orthologs mediated cell-cell fusion in each of the cell lines tested, with the exception of the pigtail macaque ortholog in human A549 cells (Figs. 2B, S4). Fusion efficiency varied between EnvP(b)1 orthologs and cell lines (Figs. 2B, S4, S5). EnvP(b)1 from great ape species (human, chimpanzee, and gorilla) were generally the most fusogenic across all cell lines. In these assays the extent of syncytia formation did not correlate with the extent of apparent EnvP(b)1 expression or processing.

### Structure of the EnvP(b)1 inferred receptor binding domain

We determined the crystal structure of the human EnvP(b)1 iRBD. Single anomalous dispersion (SAD) from a platinum derivative was used to obtain experimental phases and ultimately produce a model (Fig. 3A). The iRBD’s core is defined by a β sandwich that shares some similarity with a V-type Ig fold, although it lacks the first β sheet at the N terminus. Computational analysis suggests that the rest of the N-terminal region of EnvP(b)1 is helical, and that it is cleaved by signal peptidase. Extensions between the β sheets form a series of loops and α-helical regions that decorate the core. There is weak sequence identity between these features and the gammaretrovirus variable regions A, B, and C (VRA, VRB, and VRC, respectively), which are believed to dictate Env-receptor interactions for gammaretroviruses (15, 16).

**Fig. 3.**
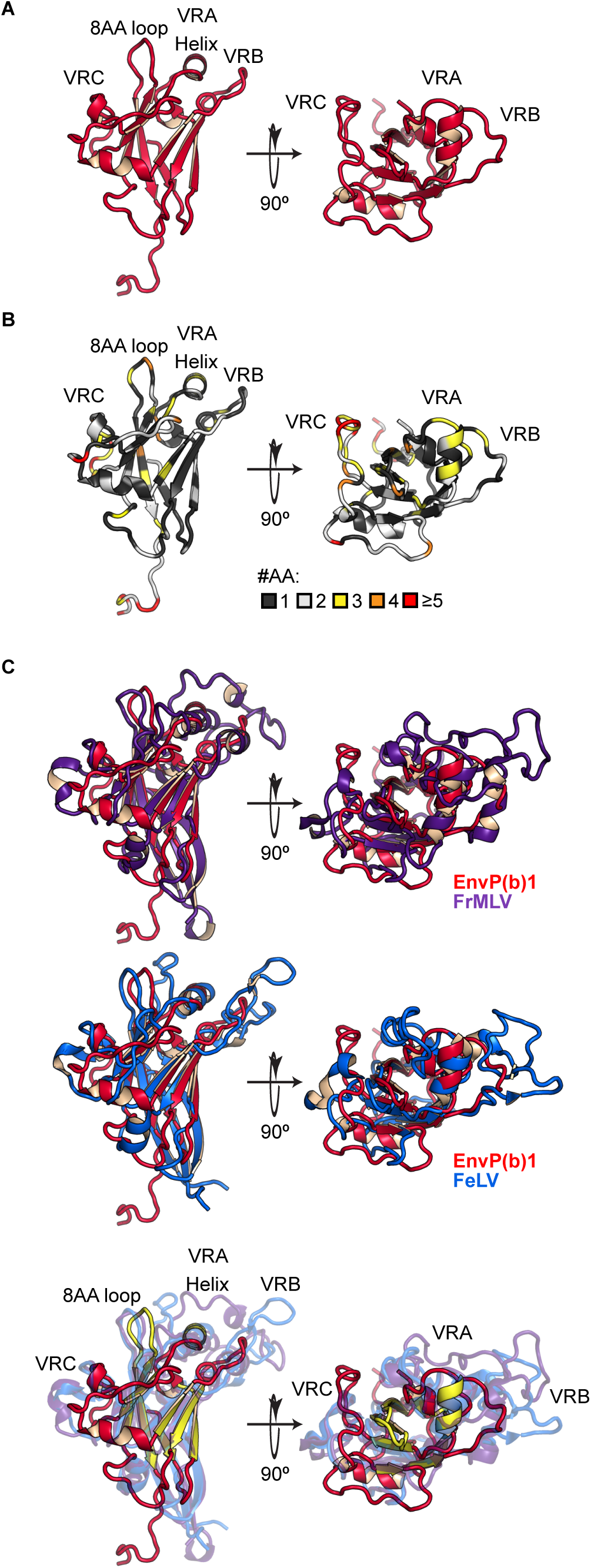
Structure of EnvP(b)1 inferred receptor binding domain (iRBD). (A) Crystal structure of EnvP(b)1 iRBD. A C-terminal tag that is present in the structure was removed in this representation. Variable regions (VRA, VRB, VRC) and the 8 amino acid loop and central helix are annotated. (B) Amino acid diversity of the simian EnvP(b)1 iRBD. Amino acid diversity was compared between 35 simian EnvP(b)1 iRBD sequences. Each residue is colored based on the number of unique amino acids found at that position. (C) Comparison of EnvP(b)1 iRBD with RBD structures of extant gammaretroviruses. Top: EnvP(b)1 (red) overlaid with FrMLV (purple) (PDB: 1AOL). Middle: EnvP(b)1 overlaid with FeLV (blue) (PDB: 1LCS). Bottom: Overlay of all three RBD structures. Common features of the beta sheet core, central helix, and 8 amino acid loop are highlighted in yellow on the EnvP(b)1 structure.

The iRBD is conserved in sequence among the 35 simian EnvP(b)1 orthologs (Fig. S3), with insertions and deletions only present in the unstructured C terminus. We mapped amino acid diversity to the iRBD structure by scoring the number of different amino acids that were present at each position (Fig. 3B). For the majority of the domain, only one or two amino acids are present at a given site. Substitutions within the core were often conservative. Most of the sites that tolerate amino acid diversity are in loops or surface exposed residues on helices. In particular, the sequence of VRC has diverged the most rapidly, suggesting that its sequence is not subject to the same constraints as other regions of the iRBD.

We compared the iRBD structure to the two other known gammaretrovirus RBD structures. These are from Friend murine leukemia virus and feline leukemia virus (FrMLV and FeLV, respectively), both extant circulating viruses (15, 17). The iRBD shares little (~20%) sequence identity with these examples, yet shows extremely strong structural similarities. The EnvP(b)1 iRBD is more compact with shorter, less prominent loop structures extending from its core (Fig. 3C). All three VRs are shorter in EnvP(b)1: VRA and VRB are dramatically shorter, and the VRC, also shorter, adopts a different conformation as it is liberated from constraints by the absence of the paired Cys residues that form a disulfide in FrMLV and FeLV. Additional connecting regions also exhibit changes in length and corresponding structural differences. As a consequence, the potential receptor-interacting interfaces of EnvP(b)1 are quite different from those for FrMLV and FeLV. However, there is clearly a structurally conserved architecture that is common among gammaretroviruses. The β sandwich core, a central helix that runs perpendicular to it, and an 8-amino acid loop between the last two β sheets are all structurally conserved. This conserved scaffold structure serves as a platform for the presentation of unique elements that determine receptor binding specificity.

## DISCUSSION

Captured viral diversity has produced host genetic novelty. Viral Env proteins have at least two activities: receptor binding and membrane fusion. In cells, Env-receptor interactions can result in receptor blockade or down-regulation of the receptor from the cell surface. Receptor engagement is also often a trigger that potentiates membrane fusion. Over the duration of animal evolution these viral functions have been coopted on multiple instances to perform host functions. Those ERV Envs that have been coopted are diverse in their ages, sequences, and, where known, binding partners (18). It appears that capturing and maintaining a variety of genes from divergent viruses provides evolutionary benefits.

EnvP(b)1 has all the hallmarks of a viral fusogen that has been repurposed to perform a host function. In contrast to other ERV-P(b)1 genes, the EnvP(b)1 ORF has been maintained as a coopted gene for the entirety of simian evolution, likely longer. At least one of its functions, membrane fusion, has also been retained in every simian clade. Our transfection and fusion experiments, performed in cells derived from divergent primate species, indicate that the unidentified cellular factors required for fusion, such as receptors and proteases, are also conserved. In these assays, ape and New World monkey EnvP(b)1 orthologs tended to behave more similarly to each other, while Old World monkey EnvP(b)1s were overall less fusogenic. These lineage-specific differences may reflect the consequences of divergence and coevolution of EnvP(b)1 orthologs with the factors that are required for cell-fusion.

EnvP(b)1 protein is expressed in multiple healthy human tissues. In these tissues it exists in a processed, presumably fusion-competent form. We detected the fully processed SU form and no unprocessed SU-TM in all EnvP(b)1-expressing tissues. In contrast, SU-TM was the major species in transfected cells. We speculate that in tissues, expression of EnvP(b)1 and the proper furin-like protease may be coordinated. Our detection of protein was generally in agreement with the previously reported RNA abundances, with the exception of splenic tissue, where we only observe a band of a different molecular weight (5, 7). Placental tissue had the highest expression of EnvP(b)1 protein. In other tissues, expression did not vary with pregnancy or gender. The function of EnvP(b)1 is unknown, but its maintenance among simians for at least 40-47 million years, and its expression and processing in human tissues indicates that it must in some way contribute to an important physiological process, likely involving membrane fusion.

Among human examples, selective preservation or loss of furin processing has accompanied cooption. The two syncytins are activated by furin-like cleavage events and act as membrane fusogens (2). HERV-T is no longer processed, is not fusogenic and has evolved to downregulate its receptor (19). HEMO, of unknown function, is not classically processed by proteases but has evolved to be shed from cells via an alternative cleavage event (20). Others, like recently integrated HERV-K copies, may have been too recently acquired to have gained or lost functions. EnvP(b)1 has clearly evolved to retain its processing and fusogenicity.

Detection of sequences related to EnvP(b)1 and the larger ERV-P(b) provirus in the genome of the aye-aye, but not other prosimian species, was unexpected. It suggests that related viruses may be more ancient than our estimate based upon the primate phylogeny. Synteny indicates EnvP(b)1 was present in the *RIN3* locus before the divergence of New and Old World monkeys. Fossil records and genetic dating approaches estimate that the split occurred 40-47 MYA (9–11). One study, using LTR dating approaches, has estimated ERV-P(b)1 to be as old as 55 MYA (6). The lemur colonization of Madagascar, 55-75 MYA (12, 13), and their subsequent geographic and genetic isolation were concurrent with or predate the simian acquisition of EnvP(b)1. After the arrival of lemurs to Madagascar, aye-ayes promptly diverged from all other extant lemur species (Lemuriformes) (9–11, 14). There are several plausible evolutionary scenarios to explain the aye-aye ERV-P(b) related sequences (Fig. S1C). We favor one in which the ancestors of simians and lemurs were independently infected by related viruses on the continent of Africa 55-75 MYA, leading to independent endogenization events, and that those ERVs were later lost from Lemuriformes. We cannot discern if EnvP(b)1 was once present in a common ancestor of all primates due to seemingly independent deletions in the intron that harbors ERV-P(b)1 in lemur and tarsier genomes. In all scenarios, it is likely that the infectious progenitor virus and EnvP(b)1 are quite ancient.

EnvP(b)1 is separated by at least 40-47 million years of evolution from related extant retroviruses for which structural data exist. We determined a structure of the iRBD from this ancient viral element. The iRBD amino acid sequence is largely conserved among the EnvP(b)1 loci found in host genomes. Though under purifying selection, some diversity has accumulated in the variable regions. These sites evolve rapidly in circulating viruses, and their divergence in EnvP(b)1 indicates that such sites can tolerate substitution without compromising function even in the context of a coopted protein. This conservation and the lack of insertions/deletions suggest that the current iRBD may closely approximate that of the infectious progenitor virus that circulated >55MYA. Comparisons with RBD structures from two extant gammaretrovirus RBDs highlights the exquisite evolutionary plasticity of this domain. All three share a conserved architecture that depends little on sequence conservation. Upon this common scaffold, each Env has evolved a series of unique features, demonstrating the versatility of this fold as a platform for recognizing many different receptor molecules. This versatility in receptor recognition may underlie the evolutionary success of gammaretroviruses, and of the Env proteins themselves, as evidenced by their wide distribution as viral and endogenous genes.

## MATERIALS AND METHODS

### Cell lines

Human 293F cells were maintained at 37°C with 5% CO_2_ in FreeStyle 293 Expression Medium (ThermoFisher) supplemented with penicillin and streptomycin. Human 293T (ATCC CRL-3216; American Type Culture Collection, Manassas, VA), Green monkey Vero (ATCC CCL-81), and Rhesus macaque LLC-MK2 (ATCC CCL-7) were grown in Dulbecco’s modified Eagle’s medium (DMEM) supplemented with 10% fetal bovine serum (FBS) and maintained at 37ºC and 5% CO_2_. Owl monkey OMK cells (ATCC CRL-1556) were grown in Eagle’s minimum essential medium (EMEM) supplemented with 10% FBS and maintained at 37ºC and 5% CO_2_.

### Identification of EnvP(b)1 sequences

Human, chimpanzee, white-cheeked gibbon, African green monkey, and pigtail macaque sequences were previously published (5). All other full length EnvP(b)1 ORFs were identified through querying publicly available databases including NCBI genbank, Ensmbl, and UCSC genome browser, using the full-length human EnvP(b)1 nucleotide sequence. Databases queried included fully assembled genomes, NCBI nucleotide database, and whole genome shotgun reads identified through the basic local alignment tool (BLAST). Details of these sequences are shown in Table S2.

Sequences mapping to the ERV-P(b) provirus in the aye-aye genome were identified through BLAST using either the SU domain of human EnvP(b)1 or the entire human ERV-P(b)1 locus, querying whole genome shotgun sequences from *Daubentonia madagascariensis* (aye-aye). 11 reads were identified that mapped to ERV-P(b) from an unassembled aye-aye genome (21). Accession numbers for these reads are: AGTM011455045, AGTM012725663, AGTM010332352.1, AGTM011595709.1, AGTM012840819.1, AGTM011264183.1, AGTM012908686, AGTM010255092, AGTM010981769, AGTM011358766, and AGTM010685344.

### Plasmids

Human EnvP(b)1 codon-optimized sequence was ordered from GenScript. EnvP(b)1 sequences from chimpanzee, white-cheeked gibbon, gorilla, African green monkey, pigtail macaque, crab-eating macaque, sooty mangabey, baboon, common marmoset, and owl monkey, were ordered from Genewiz. All sequences were cloned into a modified version of pVRC-8400 expression vector (22). EnvP(b)1^CS-^ with the furin cleavage site mutated from RKTR to SKTR was generated from the human EnvP(b)1 plasmid using QuikChange site-directed mutagenesis (Agilent). EnvP(b)1 iRBD sequences were synthesized and cloned into a modified pVRC8400 expression vector, as previously described (23, 24).

### Recombinant EnvP(b)1 iRBD expression and purification

The inferred receptor binding domain of human EnvP(b)1 was deduced as follows. A region of homology to murine and feline leukemia virus RBDs was identified using the Phyre2 server. The signal peptide cleavage site was predicted using the SignalP-4.1 Server (25), and Phyre2 secondary structure prediction (26). This defined the iRBD N terminus. The C-terminal boundary was defined by the transition from a predicted unstructured region to the proline-rich region. iRBDs were synthesized and cloned into a modified pVRC8400 expression vector, as previously described. We produced a variant with a Human Rhinovirus 3C protease cleavage site and 6xHis tag, and a variant with an HA-tag. iRBD variants were produced by polyethylenimine (PEI) facilitated, transient transfection of 293F cells that were maintained in FreeStyle 293 Expression Medium. Transfection complexes were prepared in Opti-MEM and added to cells. Supernatants were harvested 4-5 days post-transfection and clarified by low-speed centrifugation. iRBDs were purified by passage over Co-NTA agarose (Clontech) followed by gel filtration chromatography on Superdex 200 (GE Healthcare) in 10 mM Tris-HCl, 150 mM NaCl at pH 7.5 (buffer A).

For immunization of rabbits and production of antisera enriched for iRBD directed IgGs, we used the version with the 3C protease site and 6xHis tag. Both were removed by treatment with PreScission protease (MolBioTech; ThermoScientific). The protein was repurified on Co-NTA agarose followed by gel filtration chromatography on Superdex 200 (GE Healthcare) in buffer A to remove the protease, tag, and uncleaved protein. Crystallographic studies used the HA-tagged version.

### Generation and enrichment of EnvP(b)1 iRBD specific antisera

Bulk anti-iRBD serum was generated by Covance. Purified recombinant EnvP(b)1 iRBD was used to immunize a single rabbit. Serum from the terminal bleed was tested for detection of EnvP(b)1 by western blot. Crude rabbit serum was diluted 1:50 in buffer A. Bulk IgGs were captured by protein A agarose resin (GoldBio) at 4°C rotating overnight. The resin was collected in a chromatography column, washed with a column volume of buffer A and eluted with 0.1 M glycine buffer (pH 2.5) which was immediately neutralized by 1 M tris(hydroxymethyl)aminomethane (pH 8.0). Antibodies were then dialyzed against phosphate buffered saline (PBS) pH 7.4. To produce an EnvP(b)1 iRBD affinity resin, EnvP(b)1 was expressed and purified as above. Cleaved iRBD was then concentrated, dialyzed against phosphate buffered saline (PBS) pH 7.4, and arrested to Pierce™ NHS-activated agarose per manufacturer’s protocol. The reaction was then quenched by the addition of tris(hydroxymethyl)aminomethane (pH 7.5). Rabbit antibodies were then incubated with the iRBD resin at 4°C rotating overnight. The iRBD resin was collected in a chromatography column, washed with a column volume of PBS (pH7.4) and eluted with 0.1M glycine buffer (pH 2.5) which was immediately neutralized by 1M tris(hydroxymethyl)aminomethane (pH 8.0). Antibodies were then dialyzed against phosphate buffered saline (PBS) pH 7.4.

### Crystallization, structure determination, and refinement

EnvP(b)1 iRBD was concentrated to 10-12mg/ml. Crystals were grown in hanging drops over a reservoir of 0.5 M sodium chloride, 0.1 M bis-tris(hydroxymethyl)aminomethane propane pH 7.0, 20% (w/v) polyethylene glycol 4000. Platinum was incorporated by incubating crystals in a solution of 0.08 M bis-tris(hydroxymethyl)aminomethane propane, 0.4M sodium formate, 16% (w/v) polyethylene glycol 4000 and 10mM Potassium tetrachloroplatinate(II) for approximately 48 h. All crystals were cryoprotected in 22-25% (v/v) glycerol. We recorded diffraction data at the Advanced Light Source on beamline 8.2.2 and on the Advanced Photon Source on beamline 24-ID-C.

Using XDS (27), data were processed for one native and the single anomalous dispersion (SAD) data from one platinum derivative crystal. The merged data used for phasing had anomalous signal to about 3.5 Å resolution. Using SHELX (28) we found 5 platinum sites, refined them, and calculated phase probabilities. Initial maps were obtained after statistical density modification using RESOLVE (29). The Matthew coefficient suggested that there were multiple iRBD copies in the asymmetric unit. We flooded the initial map with dummy atoms and carved out a single copy of iRBD (the one with the best-resolved density) by inspection in PyMol (Schrödinger, LLC). With this we obtained a search model to locate two additional copies of iRBD in the asymmetry unit by molecular replacement using PHASER (30). After further optimizing the phases by 3-fold NCS averaging we obtained a map of reasonable quality to build an initial model that was then used to phase the native dataset.

We carried out refinement calculations with PHENIX (31) and model modifications with COOT (32). Refinement of atomic positions and B factors was followed by translation-liberation-screw (TLS) parameterization and placement of water molecules. Final coordinates were validated with the MolProbity server (33). Data collection and refinement statistics are in Table S3. Figures were made with PyMol.

### Human tissue lysates and cell lysate western blots

Tissue protein lysates were purchased from Takara. Details for each lysate are shown in Table S1. Tissue lysate was mixed with Laemmli buffer with 2-mercaptoethanol and boiled for 5 mi. 40 µg tissue lysate was run on a 4-20% acrylamide gel (Biorad).

Human EnvP(b)1, EnvP(b)1^cs-^, HERV-W Env, and empty vector were transfected into 293T cells using Lipofectamine 3000 reagent (Life technologies). Human, chimpanzee, gorilla, gibbon, African green monkey, pigtail macaque, sooty mangabey, baboon, crab-eating macaque, owl monkey, and marmoset EnvP(b)1 plasmids were transfected into A549, Vero, LLC-MK2, and OMK cells using Lipofectamine 3000 reagent. 24 h post-transfection, cells were lysed in TNE buffer (50mM Tris, 2mM EDTA, 150mM NaCl) supplemented with 1% tritonX-100. Lysates were boiled in Laemmli buffer and run on 4-20% acrylamide gels.

For all western blots, proteins were transferred onto nitrocellulose membrane. Membranes were blocked with 5% milk in PBS+0.1%Tween-20 and probed with anti-EnvP sera or mouse anti-GAPDH antibody (Genscript), followed by goat anti-mouse HRP (Invitrogen) or goat anti-rabbit HRP (Sigma-Aldrich) antibodies. Membranes were incubated with ECL reagent (Thermo) and signal was detected using an Amersham imager 600 device (GE healthcare). Images were processed using ImageJ software (National Institutes of Health).

### Cell-cell fusion

Human, chimpanzee, gorilla, gibbon, African green monkey, pigtail macaque, sooty mangabey, baboon, crab-eating macaque, owl monkey, and marmoset EnvP(b)1 plasmids were transfected into A549, Vero, LLC-MK2, and OMK cells using lipofectamine 3000 reagent. 18-24 h post-transfection, cells were stained with NucBlue live cell nuclear stain (Life technologies) for 20 minutes at 37ºC and subsequently imaged. Images were processed using ImageJ software.

## DATA AVAILABILITY

Coordinates and diffraction data have been deposited at the Protein Data Bank (PDB) (ID code: 6W5Y).

## ACKNOWLEDGEMENTS

We thank A. Kirmaier for helpful comments on this manuscript. X-ray diffraction data were recorded at beamline ID-24-C (operated by the Northeast Collaborative Access Team [NE-CAT]) at the Advanced Photon Source (APS; Argonne National Laboratory). The NE-CAT is funded by NIH grant P41 GM103403. The APS is operated for the DOE Office of Science by Argonne National Laboratory under contract DE-AC02-06CH11357 and at beamline 8.2.2 (operated by the Berkeley Center for Structural Biology and the Howard Hughes Medical Institute) at the Advanced Light Source (Lawrence Berkeley Laboratory). We thank all beamline staff members for advice and assistance in data collection. This study was funded by NIH grants AI109740 and AI059371 to S.P.J.W.

---

**Fig. S1.**
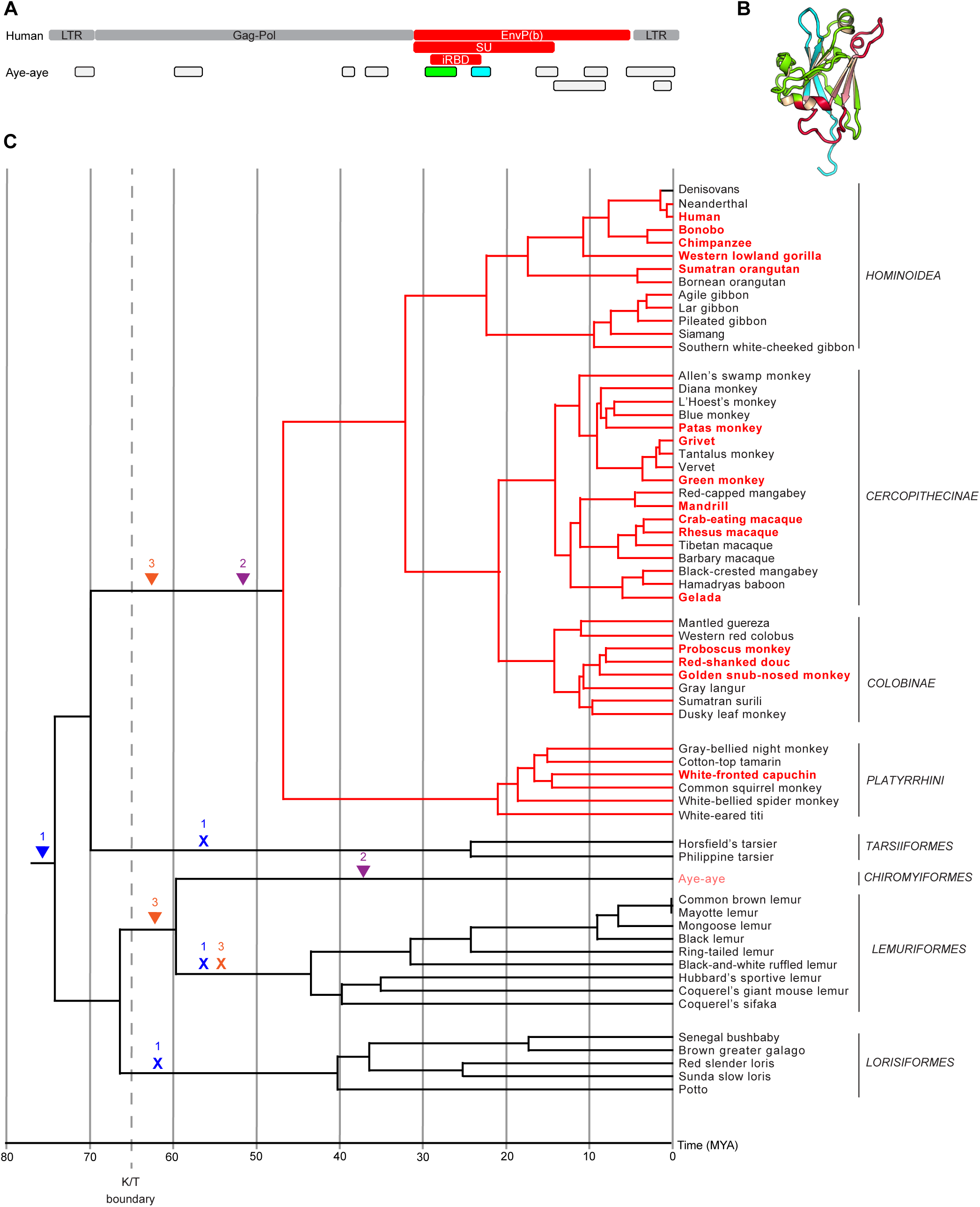
Evidence of EnvP(b) in the aye-aye genome. (A) Reads from deposited aye-aye whole genome shotgun sequencing reads map to multiple regions in ERV-P(b). Whole genome shotgun reads (Genbank accession numbers AGTM011455045, AGTM012725663, AGTM010332352.1, AGTM011595709.1, AGTM012840819.1, AGTM011264183.1, AGTM012908686, AGTM010255092, AGTM010981769, AGTM011358766, AGTM010685344) were aligned to human ERV-P(b) in the RIN3 locus. The regions to which the aye-aye sequences align are indicated. Green and cyan segments map to the EnvP(b)1 iRBD. (B) Aye-aye sequences mapped onto human EnvP(b)1 iRBD structure. Sequences in green and cyan correspond to those in panel A. (C) Proposed scenarios for ERV-P(b) integration into the germline of simians and aye-ayes. Arrowheads indicate integration events. X indicates loss. 1. Integration into the last common ancestor of all primates, followed by loss in the Lorisiformes, Lemuriformes, and Tarsiiformes lineages. 2. Asynchronous integration into the last common ancestor of Simiiformes on mainland Africa and into the Chiromyiformes on Madagascar. 3. Independent integrations into multiple primates on Africa prior to lemur colonization of Madagascar followed by loss in the Lemuriformes. Tree adapted from Pozzi et al., 2014 (9). Species highlighted in red are those in which we identified an intact EnvP(b)1 ORF. Branches highlighted in red indicate lineages in which ERV-P(b) insertion is unambiguous, in which we identified EnvP(b)1 in sister species not included in this tree. Aye-aye is highlighted in pink.

**Fig. S2.**
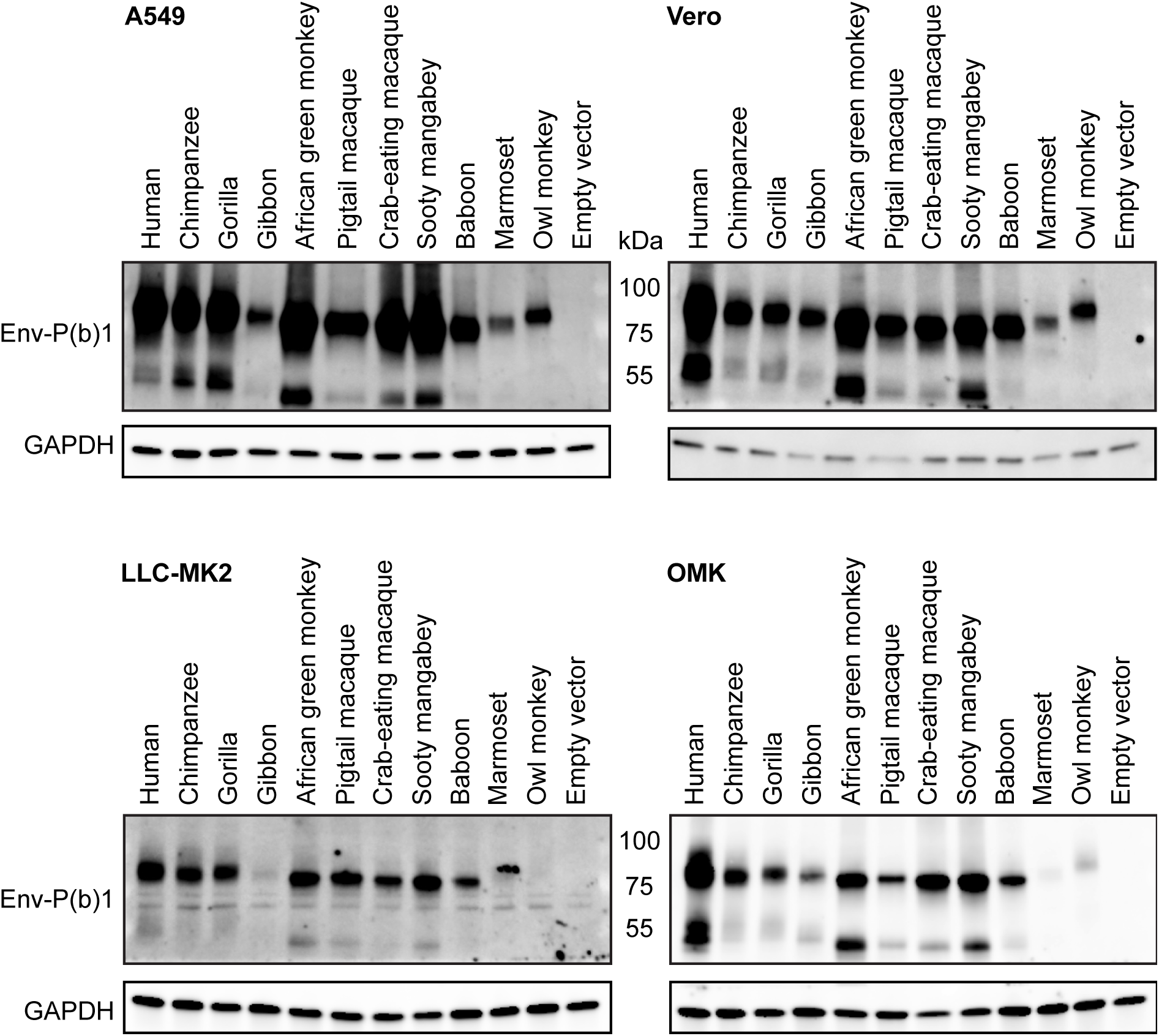
Expression of EnvP(b)1 from multiple primates in primate cell lines. Cell lysates from the indicated cells lines transfected with the EnvP(b)1 from the indicated species were subjected to western blotting with either anti-EnvP(b)1 sera or anti-GAPDH antibody.

**Fig. S3.**
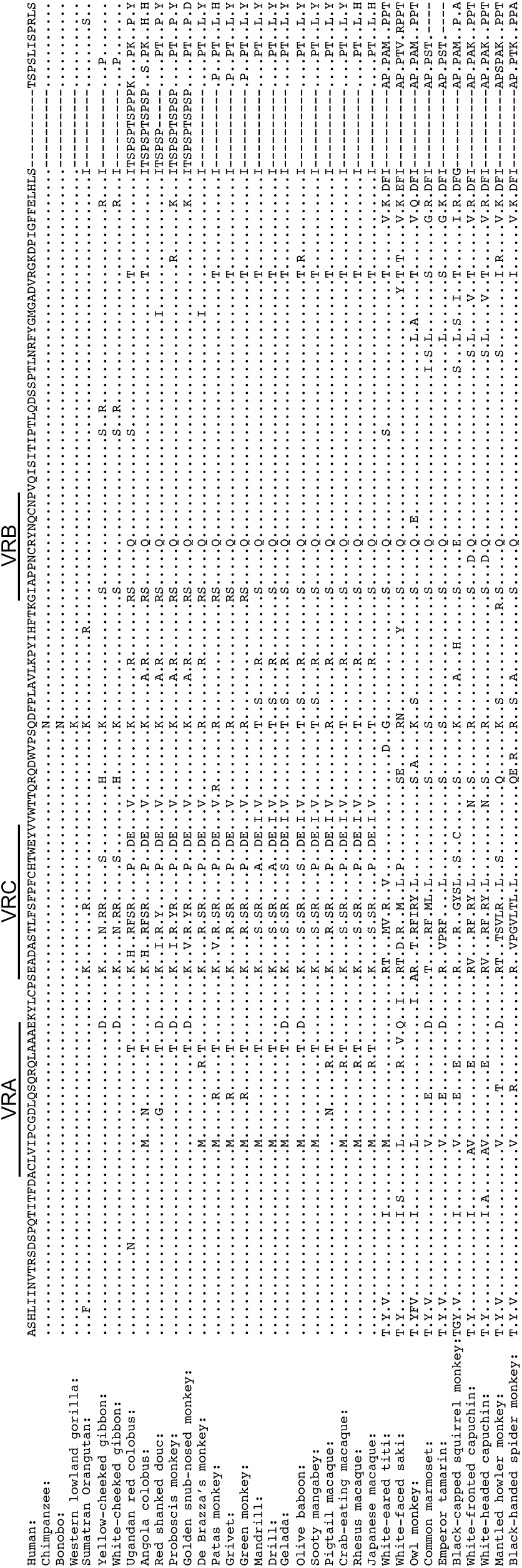
Alignment of iRBD amino acid sequence from 35 simian species. Variable regions VRA, VRB, and VRC are annotated. This alignment was used to determine the amino acid variability at each site, shown in Fig. 3B.

**Fig. S4.**
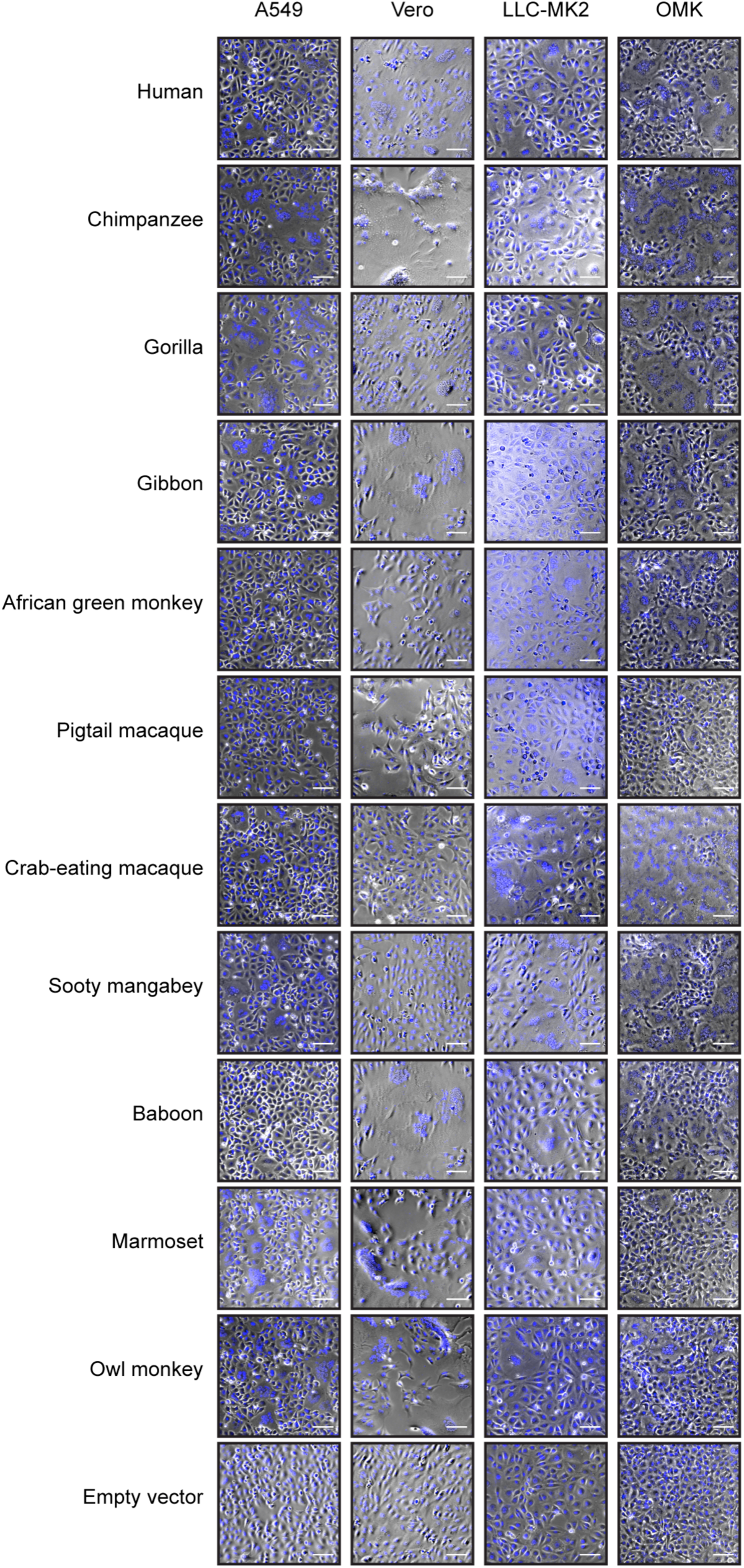
Preservation of fusogenicity of EnvP(b)1. (A) Examples of cell fusion results. The indicated cell lines were transfected with EnvP(b)1 ORFs from the indicated species. Cells were stained with nuclear stain (blue) and imaged. Scale bar: 100 µm. Cell line species: A549 – human, vero – green monkey, LLC-MK2 – macaque, OMK – owl monkey.

**Fig. S5.**
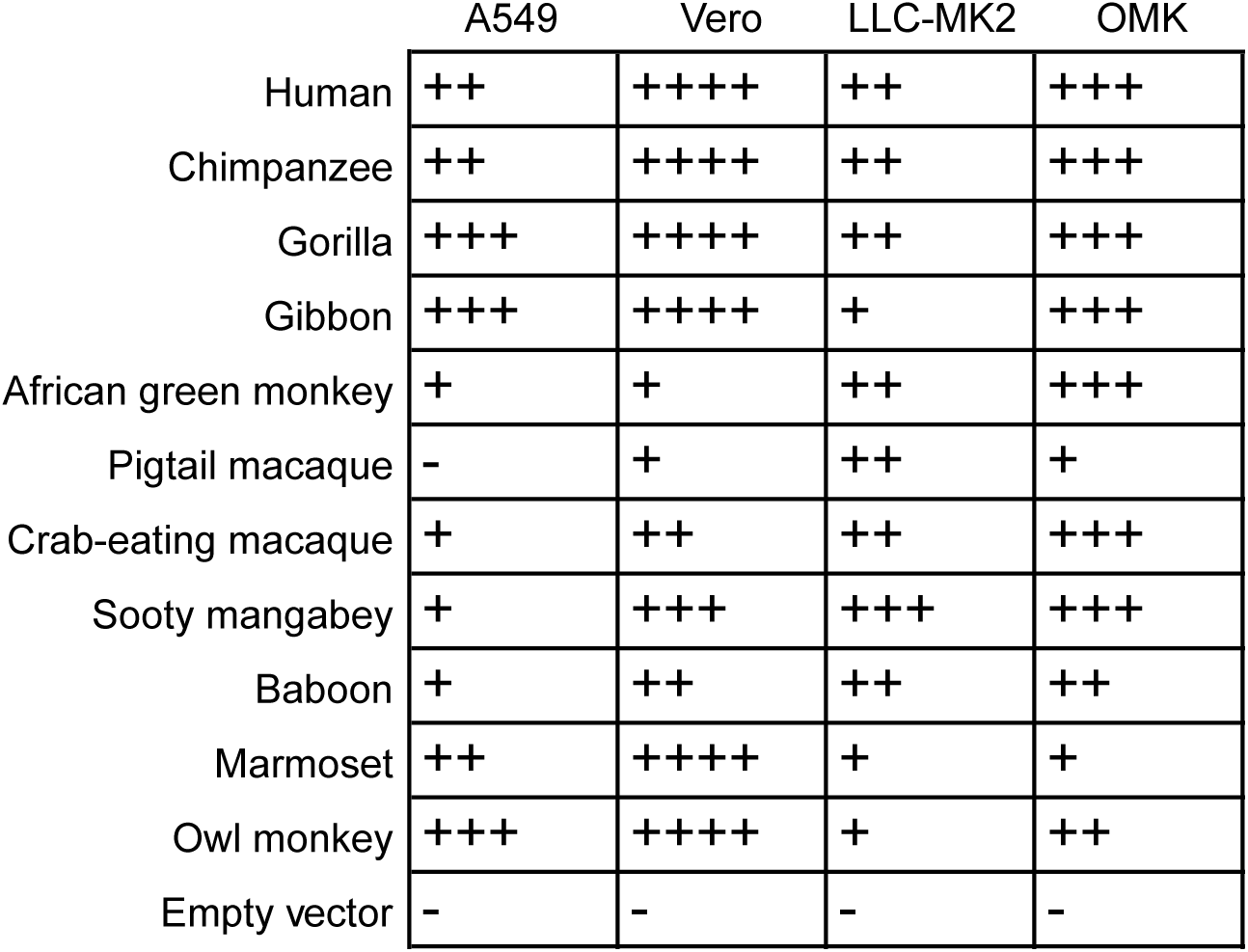
Summary of fusion data for all tested EnvP(b)1 sequences. Fusogenicity of each EnvP(b)1 in each cell was determined by approximating the percent of total cells fused in each condition. +: 1-25 ++:26-50 +++: 51-75 ++++: 75-100. Cell line species: A549 – human, vero – green monkey, LLC-MK2 – macaque, OMK – owl monkey.

**Table S1.** Information on the human tissue lysates probed for EnvP(b)1 expression. Tissue protein lysates were purchased from Takara. Some lysates were from a single donor and some were pooled from multiple donors. Information on the age, sex, and cause of death (COD) of the donors is presented. Donors were presumed to be healthy prior to death.

| Tissue | Tissue condition | #of donors | Sex | Age | COD | Cat # | Lot # |
| --- | --- | --- | --- | --- | --- | --- | --- |
| Placenta | Normal, whole placenta | 15 | F | 19-33 |  | 635307 | 1510952A |
| Testis | Normal, whole testes | 18 | M | 19-64 | Truama | 635309 | 1602008A |
| Ovary | Normal, whole ovary | 1 | F | 30 | Trauma | 635308 | 1510953A |
| Spleen | Normal, whole spleen | 1 | M | 55 | Sudden death | 635312 | 1404375A |
| Thymus | Normal | 1 | M | 21 | Sudden death | 635350 | 1002151A |
| Small Intestine | Normal, whole small intestine | 9 | M&F | 21-45 | Sudden death, trauma | 635311 | 1205370A |

**Table S2.**
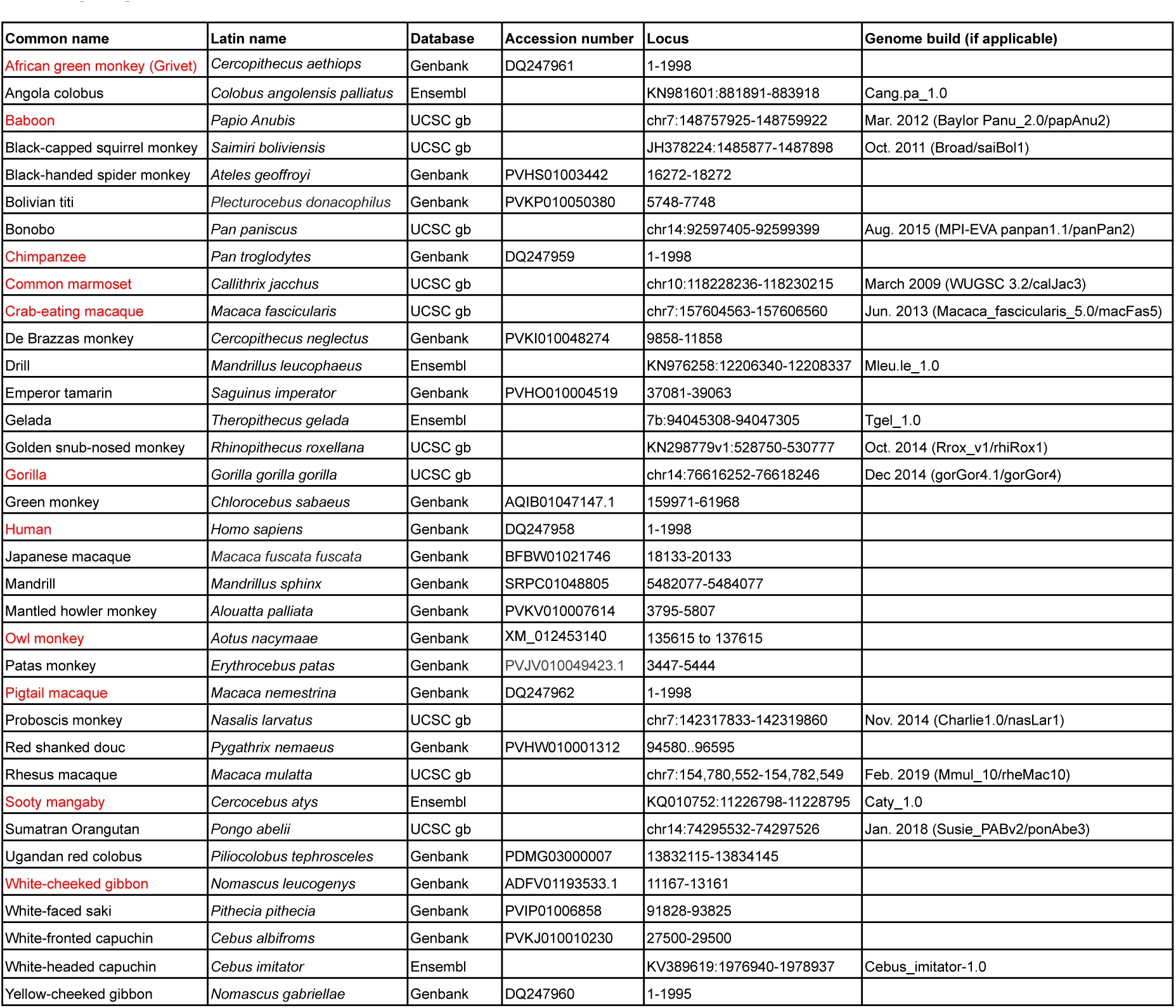
Sequence information for simian EnvP(b)1 ORFs. Loci reference nucleotide sequence. EnvP(b)1 from species highlighted in red were used for expression and fusion experiments.

**Table S3.**
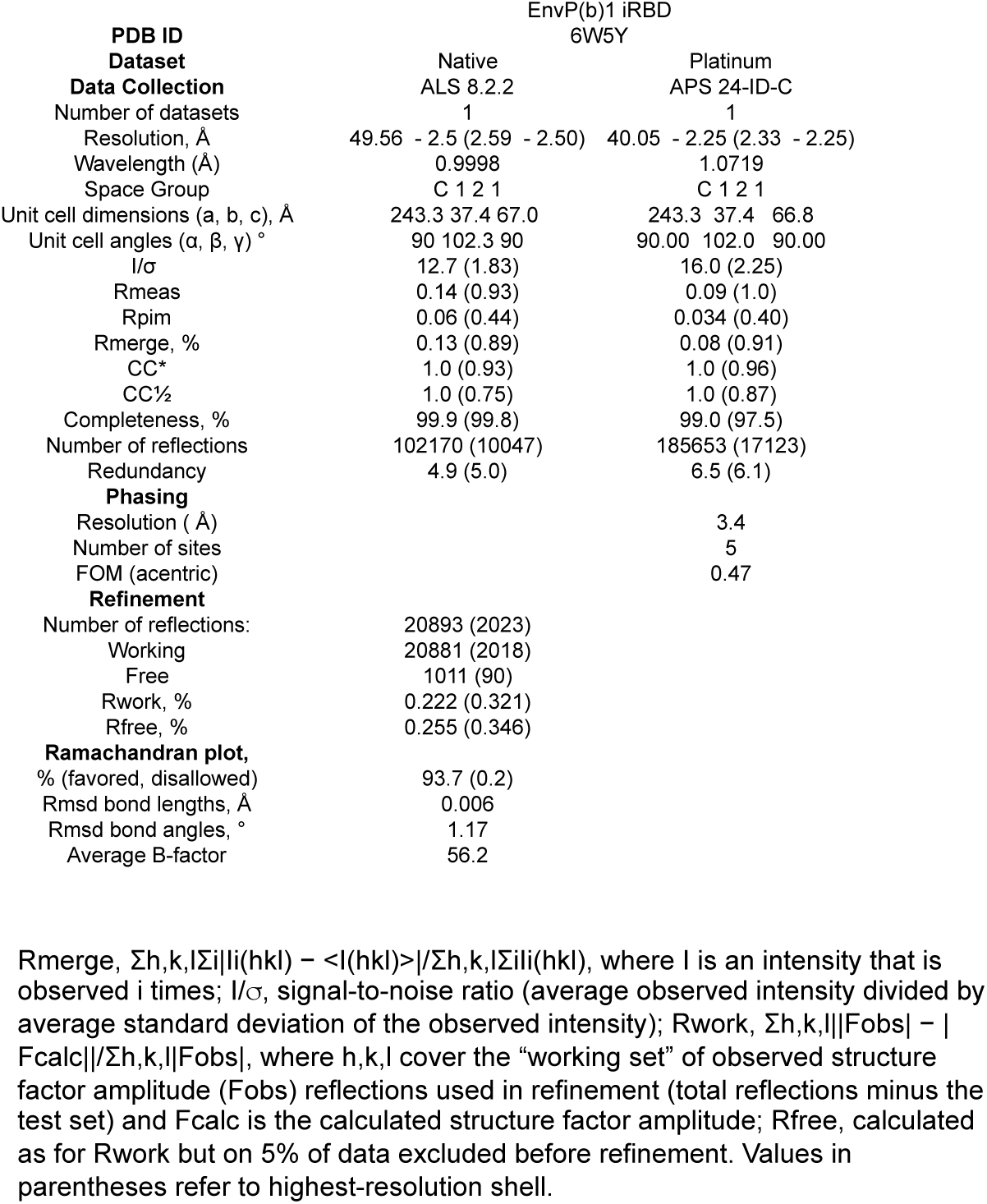
Crystallographic statistics for EnvP(b)1 iRBD

